# *Keetia magassoubiana sp*. *nov*. (Rubiaceae - Vanguerieae) a threatened evergreen forest climber of West Africa

**DOI:** 10.1101/2024.06.21.600008

**Authors:** Martin Cheek, Joel Bowden-Pickstock

## Abstract

*Keetia magassoubiana* Cheek, an evergreen rainforest climber, is described and illustrated from the Republic of Guinea, Sierra Leone, Liberia and Ivory Coast. Previously indicated as being close to but different from *Keetia tenuiflora* (Hiern) Bridson, it differs in the glossy black, glabrous, epidermis of the distal stem internodes, the first internode rarely with very sparse red adpressed hairs (vs epidermis pale white-brown, dense pale yellow spreading hairs) and abaxial leaf surface with domed domatia with a central aperture (vs domatia absent or obscure), the secondary stem leaf bases acute (vs obtuse to truncate), the bracts forming a laciniate sheath on the distal peduncle (vs two opposite triangular bracts), the pyrene surface honeycombed with pits (vs entire). The species was earlier included within *Canthium multiflorum* (Schum. & Thonn.) Hiern (now *K. multiflora* (Schum. & Thonn.) Bridson) in the Flora of West Tropical Africa second edition. An updated key is presented to the 16 species of the genus from West Africa.

*Keetia magassoubiana* is provisionally assessed using the IUCN standard as Endangered EN B1ab(iii) due to only five of the recorded 14 locations having extant forest habitat, and because of ongoing threats of habitat clearance mainly for agriculture, but also mining.

## Introduction

*Keetia* E. Phillips (Rubiaceae, Vanguerieae) was segregated from *Canthium* Lam. by Bridson (1985, 1986). This genus of about 41 accepted species (Cheek & Onana 2024) is restricted to sub-Saharan Africa (excluding Madagascar and the Mascarene Islands) and extends from Senegal and Guinea in West Africa (Gosline *et al*. 2023a; 2023b) to Sudan (Darbyshire *et al*. 2015) in the North and East, and S. Africa in the South (Bridson 1986). *Keetia* differs from other African genera of Vanguerieae by its pyrenes with a fully or partly-defined lid-like area around a central crest, and endosperm with tanniniferous areas (Bridson 1986). *Keetia* species are usually climbers (very rarely shrubby) and occur mostly in forest habitats, sometimes in wooded grassland. In a phylogenetic analysis of the tribe based on morphology, nuclear ribosomal ITS and chloroplast *trnT-F* sequences, Lantz & Bremer (2004), found that based on a sample of four species, *Keetia* was monophyletic and sister to *Afrocanthium* (Bridson) Lantz & B. Bremer with strong support. Highest species diversity of *Keetia* is found in Cameroon and Tanzania, both of which have about 15 taxa (Onana 2011; POWO, continuously updated) and in Gabon, where 10 species are currently recorded (Sosef *et al*. 2006) but around 25 are actually present, many of them undescribed (Lachenaud pers. comm. to Cheek, 2024). Several *Keetia* species are point endemics, or rare national endemics, and have been prioritized for conservation (e.g. Onana & Cheek 2011; Couch *et al*. 2019; Murphy *et al*. 2023) and one threatened species, *Keetia susu* Cheek has a dedicated conservation action plan (Couch *et al*. 2022).

Bridson’s (1986) account of *Keetia* was preparatory to treatments of the Vanguerieae for the Flora of Tropical East Africa (Bridson & Verdcourt 1991) and Flora Zambesiaca (Bridson 1998). Pressed to deliver these, she stated that she could not dedicate sufficient time to a comprehensive revision of the species of *Keetia* outside these areas: “full revision of *Keetia* for the whole of Africa was not possible because the large number of taxa involved in West Africa, the Congo basin and Angola and the complex nature of some species would have caused an unacceptable delay in completion of some of the above Floras. […] A large number of new species remain to be described.” (Bridson 1986). Several of these new species were indicated by Bridson (1986), and other new species by her arrangement of specimens in folders that she annotated in the Kew Herbarium. One of these species was later taken up and published by Jongkind (2002) as *Keetia bridsoniae* Jongkind. In the same paper, Jongkind discovered and published *Keetia obovata* Jongkind based on material not seen by Bridson. Based mainly on new material, additional new species of *Keetia* have been published by Bridson & Robbrecht (1993), Bridson (1994), Cheek (2006), Lachenaud *et al*. (2017), Cheek *et al*. (2018a), Cheek & Bridson (2019), Cheek & Onana (2024), Cheek & Bissiengou (2024), Cheek *et al*. (2024a) and there are several other specimens that fit no other species, (e.g.Cheek *et al*. 2004; 2011) and remain to be described.

In this paper we continue the project towards an updated taxonomic account of *Keetia* by describing a further new species, *K. magassoubiana* Cheek. This taxon was first indicated by Bridson (1986) who referred to it as *Keetia sp. aff. tenuiflora* (Bridson 1986:992). Under *Keetia tenuiflora* (Hiern) Bridson, she stated: “The following specimens from Sierra Leone - *Scott Elliot* 4503 & *Thomas* 6944 & 7068 - were cited by Hepper in Fl. W. Trop. Afr. Ed. 2, 2: 182 (1963) under *C. multiflorum* (Schum. & Thonn.) Hiern. These, together with more recent collections from Mali, Liberia and the Ivory Coast, are very close to *K. tenuiflora*, but probably represent a distinct infraspecific taxon. This taxon has glabrous stems with dark-coloured bark, while typical *K. tenuiflora* has stems with yellowish adpressed hairs and buff-coloured bark.” We confirm these results, have found additional characters separating these two taxa which justify species rather than subspecies rank for the new taxon (seeTable 1). We were not able to find material from Mali and suspect this refers to a specimen of the taxon from Guinea with a misleading label annotation. Recent collecting has extended the range of the taxon through much of Guinea.

**Table 1.**
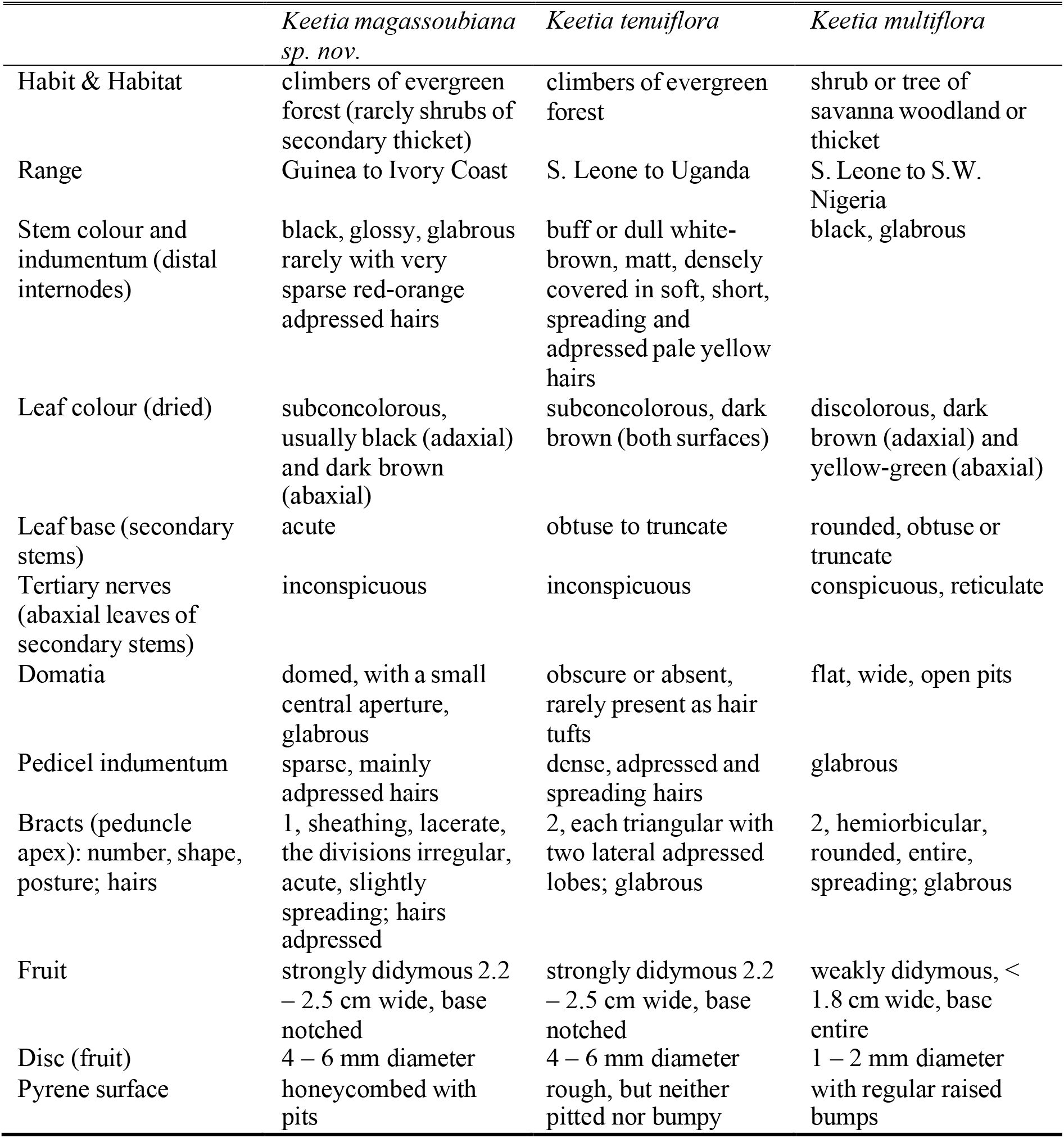
Characters distinguishing *Keetia magassoubiana sp. nov*. from *Keetia tenuiflora and Keetia multiflora*.

|  | *Keetia magassoubiana*<br>sp. nov. | *Keetia tenuiflora* | *Keetia multiflora* |
| --- | --- | --- | --- |
| Habit & Habitat | climbers of evergreen forest (rarely shrubs of secondary thicket) | climbers of evergreen forest | shrub or tree of savanna woodland or thicket |
| Range | Guinea to Ivory Coast | S. Leone to Uganda | S. Leone to S.W. Nigeria |
| Stem colour and indumentum (distal internodes) | black, glossy, glabrous rarely with very sparse red-orange adpressed hairs | buff or dull white-brown, matt, densely covered in soft, short, spreading and adpressed pale yellow hairs | black, glabrous |
| Leaf colour (dried) | subconcolorous, usually black (adaxial) and dark brown (abaxial) | subconcolorous, dark brown (both surfaces) | discolorous, dark brown (adaxial) and yellow-green (abaxial) |
| Leaf base (secondary stems) | acute | obtuse to truncate | rounded, obtuse or truncate |
| Tertiary nerves (abaxial leaves of secondary stems) | inconspicuous | inconspicuous | conspicuous, reticulate |
| Domatia | domed, with a small central aperture, glabrous | obscure or absent, rarely present as hair tufts | flat, wide, open pits |
| Pedicel indumentum | sparse, mainly adpressed hairs | dense, adpressed and spreading hairs | glabrous |
| Bracts (peduncle apex): number, shape, posture; hairs | 1, sheathing, lacerate, the divisions irregular, acute, slightly spreading; hairs adpressed | 2, each triangular with two lateral adpressed lobes; glabrous | 2, hemiorbicular, rounded, entire, spreading; glabrous |
| Fruit | strongly didymous 2.2 – 2.5 cm wide, base notched | strongly didymous 2.2 – 2.5 cm wide, base notched | weakly didymous, < 1.8 cm wide, base entire |
| Disc (fruit) | 4 – 6 mm diameter | 4 – 6 mm diameter | 1 – 2 mm diameter |
| Pyrene surface | honeycombed with pits | rough, but neither pitted nor bumpy | with regular raised bumps |

New species to science are continually being described from the range of the new species in W Africa, mainly from the surviving remnants of species-diverse forests. These include species of climbers e.g. in *Monanthotaxis* Baill. (Annonaceae,Hoekstra *et al*. 2021), *Cucumis* L. (Cucurbitaceae, Schaeffer 2021), *Hibiscus* L. (Malvaceae,Cheek *et al*. 2020a), *Keita* Cheek (Olaceae, Cheek *et al*. 2024b), small trees and shrubs e.g. *Noronhia* Thouars (Oleaceae,Jongkind 2020), *Tarenna* Gaertn. (Rubiaceae,Jongkind 2021), and *Tabernaemontana* L. (Apocynaceae,Jongkind & Lachenaud 2022), rheophytes e.g. *Ctenium* Panz., *Inversodicraea* Engl. and *Saxicolella* Engl. (Xanthos *et al*. 2021,Cheek *et al*. 2017;2022) to terrestrial herbs *Benna* Burgt & Ver.-Lib. (Melastomataceae,van der Burgt *et al*. 2022) and *Trichanthecium* Zuloaga & Morrone (Poaceae,Xanthos *et al*. 2020).

## Materials and Methods

Names of species and authors follow IPNI (continuously updated) and nomenclature follows Turland *et al*. (2018). Herbarium material was collected using the patrol method e.g.Cheek & Cable (1997). Herbarium specimens were examined with a Leica Wild M8 dissecting binocular microscope fitted with an eyepiece graticule measuring in units of 0.025 mm at maximum magnification. The drawing was made with the same equipment with a Leica 308700 camera lucida attachment. Pyrenes were characterized by simmering selected ripe fruits in water until the flesh softened and could be removed by scalpel. A toothbrush was then used to clean the pyrene surface to expose the surface sculpture and the lid. Finally, a fine saw was used to cut a transverse section of the fruit and seed, allowing observation of tanniferous cells in the endosperm and measurement of the endocarp thickness. Specimens were inspected from the following herbaria: BM, FHO, HNG, K, P, SL and YA. All Gbif.org images of specimens of *Keetia multiflora* (in which most specimens had previously been included) were checked and additional duplicates of confirmed specimens are recorded in “collections studied” below. One or two additional potential specimens of this species were also viewed on Gbif.org but image resolution was insufficient to allow confirmation (indumentum and domatia could not be seen in detail). Grid references and altitudes added posthoc, using Google Earth Pro, are indicated in brackets. They were used to assess the continued survival of the species using as proxy the continued existence of forest habitat at the collection site, and also to calculate extent of occurrence sensu IUCN (2012) for the conservation assessment. The format of the description follows those in other papers describing new species of *Keetia*, e.g.Cheek *et al*. (2024a). Terminology follows Beentje & Cheek (2003). All specimens indicated “!” have been seen. The conservation assessment follows theIUCN (2012) standard. Herbarium codes follow Index Herbariorum (Thiers, continuously updated).

### Keetia magassoubiana Cheek sp. nov

Type: Sierra Leone, Northern Region, Southern Sula Mts, Tonkolili District, Simbili Hill, western slope, 8°58’11.8”N 11°41’30.8”W, 770 m, fl. 6 Dec. 2009, *Robinson* 5 (holotype K barcode K000678056; isotypes FBC, SL).Fig. 1 &2

**Figure 1.**
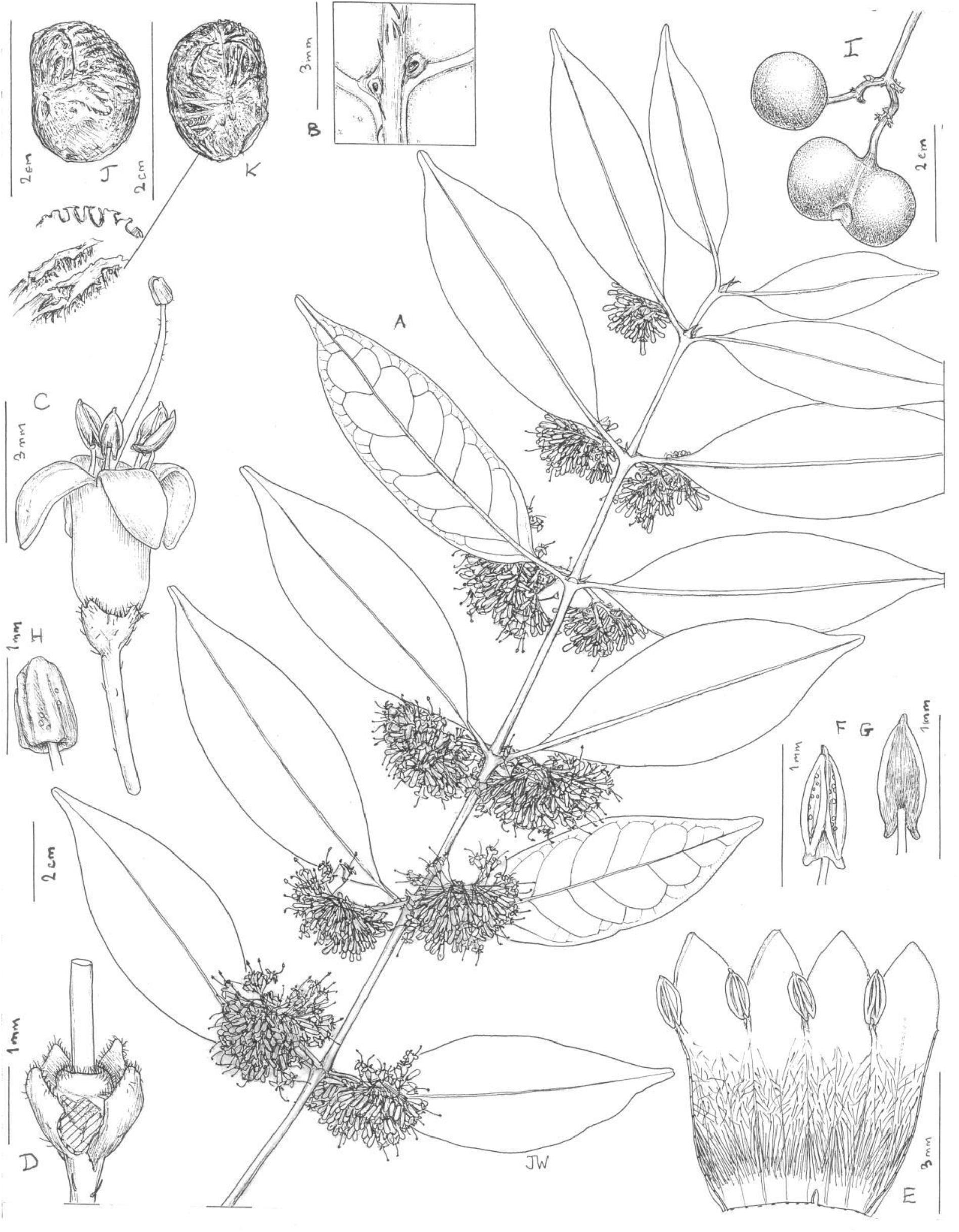
*Keetia magassoubiana* A habit, secondary, flowering branch; B detail of domed domatia and hairs, abaxial leaf surface; C flower, side view; D flower, corolla removed, calyx opened, showing disc and style; E corolla opened showing stamens and indumentum; F anther and filament, dorsal view; G as F, ventral view; H style head; I fruit both 1-seeded by abortion (top left) and 2-seeded (bottom right) showing accrescent disc; J pyrene, side view; K pyrene front view, with detail showing excavated surface. A-H from *Robinson* 5; I from *Bidault* 2434; J&K from *Jaeger* 140. Drawn by Juliet Williams.

**Figure 2.**
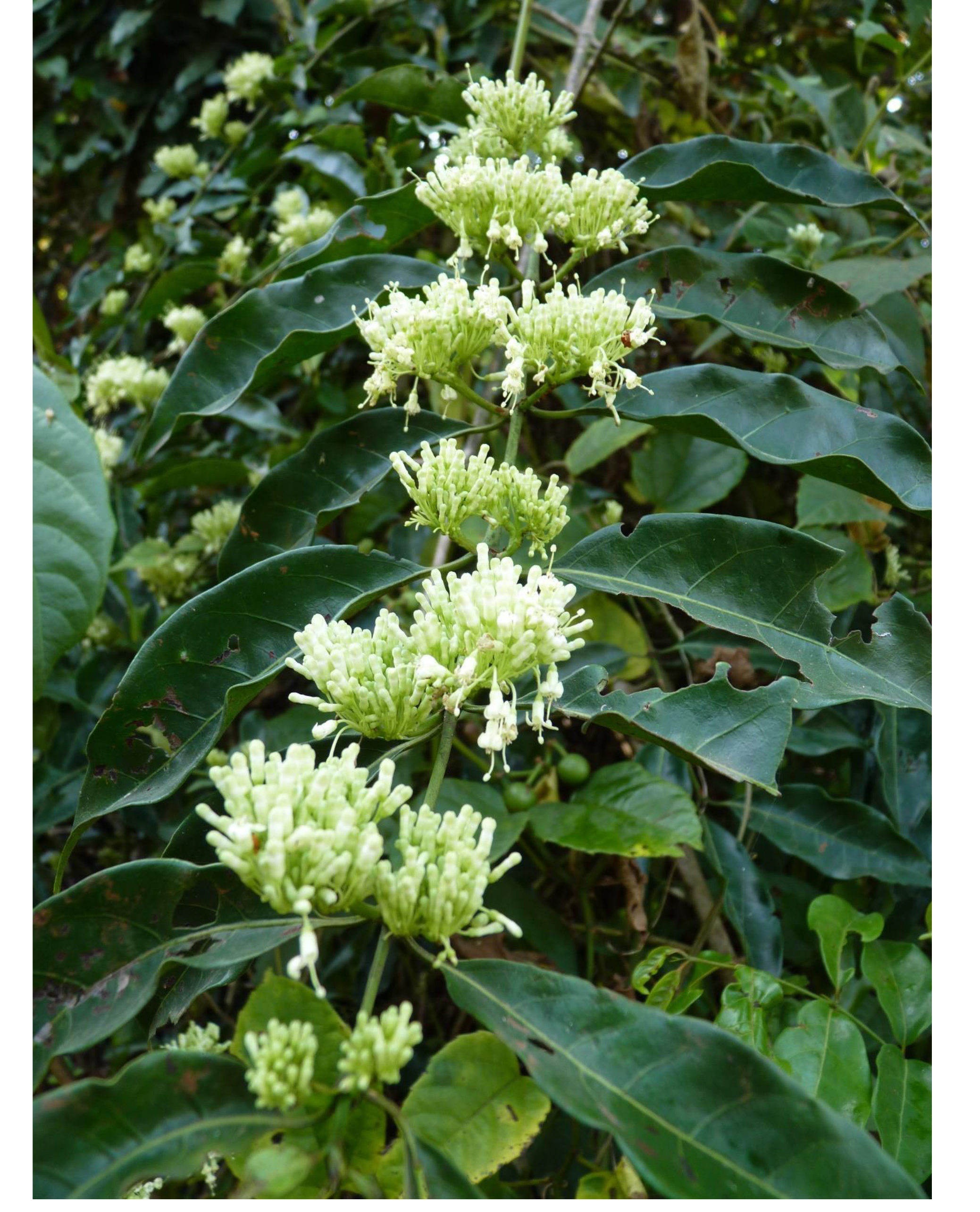
*Keetia magassoubiana* Flowering secondary branch on habitat, Sula Mts (Tonkolili) of Sierra Leone, from *Robinson* 5 (holotype K, isotypes FBC and SL). The species is now extinct at this site. Photo by Iain Darbyshire.

*Canthium multiflorum* (Schum. & Thonn.) Hiern sensu Hepper (1963: 184) pro parte quoad *Scott Elliot* 4503, *Thomas* 6944 & 7068

*Keetia sp. aff. tenuiflora* (Hiern)Bridson (1986:992).

*Keetia multiflora* (Schum. & Thonn.) Bridson sensu Aké Assi (2001:78) pro parte quoad *W. J. J. O. de Wilde* 762;Hawthorne & Jongkind (2006:624) pro parte

*Keetia tenuiflora* (Hiern)Bridson (1986:992) sensu Aké Assi (2001:78) pro parte quoad *Aké Assi* 13154 and *Aké Assi* 5321.

*Evergreen forest liana to 8 m high*, less usually a shrub of forest-derived thicket c. 3m tall. Primary (orthotropic) stem terete or rarely slightly angled, drying glossy black, at length dull white, lenticels absent, leaves, and stipules usually absent when reproductive, internodes 6 – 10 x 0.4 – 0.8 cm, glabrous. *Secondary stems* (plagiotropic, spur shoots or brachyblasts) 5 – 35 cm long, with 2 – 11 nodes, slightly ascending when young, as the primary stem, terete, 1 – 2.5 mm diameter, glabrous, or rarely the first internode with inconspicuous sparse thick dull red adpressed hairs c. 0.5mm long. *Leaves* strongly dimorphic, opposite, equal, distichous, at anthesis often present only on secondary stems, drying black on upper surface, dark brown on lower, usually leathery, bacterial nodules absent. *Leaves of primary stems* erect, held parallel to the stem, coriaceous, ovate to suborbicular 1.2 – 2.1 x 1.1 – 1.4 cm, apex rounded. Base truncate, sometimes decurrent to the top of the petiole, secondary nerves white, (1 –)2 – 3 on each side of the midrib, all arising from the proximal 1/3 – ¼ of the midrib (subpalmate), glabrous, domatia absent. Petioles canaliculate, adaxial surface broadly concave, 0.3 – 0.55(– 0.7) x 0.05 – 0.08 cm, with a few sparse slightly spreading hairs. *Leaves of secondary stems* subconcolorous, matt, adaxial surface drying black, abaxial dark brown, narrowly elliptic to oblong, (4.0 –) 5.2 – 8.8 (– 10.5) x (1.6 –) 2 – 3.8 (– 4.2) cm, shortly acuminate, acumen (0.2 –) 0.5 – 1.2 (– 1.9) cm long, apex rounded, base acute, slightly asymmetric, one side arising from the petiole 1 – 2 mm above the other; midrib and secondary nerves raised on upper surface (Fig. 1E). Secondary nerves (3 –) 4 – 6 (– 7) on each side of the midrib, nerves arising at c. 70°, straight towards margin, forking at a domatium or arching upwards towards the nerve above forming a weak, looping marginal nerve. Tertiary nerves sparse and inconspicuous, margin slightly revolute. Domatia axillary between midrib and secondary nerves and at divisions of the secondary nerves, sporadic, domed, 0.5 – 0.75mm diam, glabrous, with a minute central aperture c.0.1 diam. (Fig. 1B); indumentum restricted to the abaxial midrib, sparse, hairs reddish orange or red, adpressed, stout, straight, c. 0.5 mm long, acute; petiole of secondary stems canaliculate with the adaxial face grooved, (0.4 –) 0.6 – 1.2 (– 1.9) x 0.06 – 0.09 cm. indumentum as midrib. *Stipules* moderately persistent on secondary stems, 3.2 – 7 mm long, very shortly sheathing, base broadly triangular 1.2 – 2 x (1.8 –) 2.5 – 3 mm, midrib raised, extended as a cylindrical awn, awn straight, usually elevated and displaced to one side, (3.5 –) 4 – 4.5 x 0.25 – 0.4 mm, indumentum of the outer surface as the secondary shoots, inner surface glabrous, apart from a line of colleters and hairs at the base, sometimes exposed if the whole stipule falls; colleters 5 – 9 per stipule, drying black, glossy, stout, abaxial face flat, 0.7 –0.9 x 0.2 mm, apex acute or rounded, separated from the stem by a layer of parallel long simple, yellow hairs. *Inflorescence* axillary on the secondary stems only, held erect, above the stem, in pairs at 4 – 7 successive nodes of leafy stems, each (38 –) 50 – 110-flowered; peduncle (0.25 –) 0.3 – 0.55 cm long, glabrous; bract inserted 1 – 1.5 mm from apex, collar-like (amplexicaul, sheathing) c. 1 mm long, laciniate, lobes 6 – 7, spreading slightly, narrowly ovate-triangular, margins densely yellow-hairy; partial peduncles (branches) two, 0.2 – 0.3 cm long, subsequent branches short, bearing numerous flower fascicles, and subtending bracteoles, bracteoles alternate, ovate-lanceolate, c. 1 mm long, decreasing in size distally, indumentum as bracts. *Flowers* unpleasantly scented, of urine (Fig. 1C). Pedicels 3 – 6.5 x 0.2 – 0.3 mm, with sparse simple, yellow hairs c. 0.2 mm long, spreading, slender (Fig. 1C); calyx-hypanthium campanulate 1 – 1.5 x 1.2 – 1.5 mm, indumentum as pedicel (Fig. 1D), calyx limb tube (0.1 –) 0.3 –0.4 mm long, lobes 4 (– 5), erect, triangular 0.1 – 0.5 x 0.2 – 0.5 mm apex rounded to obtuse, indumentum on outer surface as pedicel, densest along margins, inner surface glabrous, colleters not seen (Fig. 1D). *Corolla* greenish white, in bud cylindric, c. 3.5 – 1.2 mm, apex rounded; at anthesis tube cylindrical 2.7 – 3.5 x 1.5 mm, lobes 4, about half as long as tube, reflexed, oblong-triangular, 1.6 – 2 x (1–)1.3 – 1.5 mm, outer corolla glabrous, inner surface of tube with dense, reflexed straight, simple hairs 0.9 – 1.2 mm long, originating in a band 0.5 mm wide, 1.5 mm from the base of the tube, leaving the basal c. 0.5 mm of the tube glabrous; the middle part of the tube with shorter, slightly wavy mainly downward-pointing hairs 0.2 – 0.4 mm long, obscuring the surface, reducing in density towards the throat, the distal 0.5 mm of the tube mainly glabrous, inner surface of corolla lobes glabrous (Fig. 1E). *Stamens* 4 (– 5) inserted just inside the throat, entirely exserted, erect; filaments 0.3 – 0.5 mm long, glabrous, sub-basifixed; anthers subsagittate, 0.8 – 1 x 0.3 – 0.4 mm, connective flat, brown covering the abaxial surface, extending as a mucro 0.08 mm long (Fig. 1F&G); disc torus-like 0.25 x 0.7 mm, puberulent, hairs patent, 0.8 mm long, dense (Fig. 1D). *Style* 7 x 0.25 mm, exserted for most of its length, extending beyond the anthers, distal part tapering to 0.1 mm wide, and with scattered patent hairs, 0.05 mm long; style-head cylindrical 0.6 – 0.8 x 0.5 – 0.6 mm, outer surface with c. 10 shallow longitudinal furrows (Fig. 1H). *Fruit* green when mature, surface smooth, glossy, 2-seeded fruits strongly didymous 1.5 x 2.2 – 2.5 cm, with a deep groove on both sides separating the globose carpels, emarginate at base and apex, disc bright white, flat, markedly accrescent, c. 0.5 cm diam., calyx lobes inconspicuous (Fig. 1H); unicarpellate fruits globose, 1.2 – 1.4 cm diam., disc lateral, separated from the pedicel by the vestigial carpel, vestigial carpel domed or broadly conical, c. 2 mm long, c. 3 mm diam., mesocarp c. 2 mm thick, surface glabrous. *Pyrene* broadly ellipsoid, 1.3 – 1.5 x 1.05 – 1.2 x1.05 – 1.15 cm, apex and base broadly rounded, ventral surface slightly less convex than others, point of attachment central; lid-like area entirely on ventral surface, semi-circular c.0.7 x 0.8 cm, the distal arced perimeter with a deep fissure, the proximal margin transverse, straight, poorly defined, crest and ridges absent, surface of pyrene, including lid, honey-combed with irregularly longitudinal excavations radiating from point of attachment (Fig. 1J&K); endocarp 1 – 2 mm thick. *Seed* ellipsoid, c. 7 x 5 x 5 mm, surface brain like, pale brown; endosperm (in transverse section) with slender tanniniferous bands radiating from the central embryo.

### RECOGNITION FEATURES

*Keetia magassoubiana* Cheek sp. nov. is close to but differs from *Keetia tenuiflora* (Hiern) Bridson in the glossy black, glabrous, epidermis of the distal stem internodes, rarely with very sparse red hairs (vs epidermis pale white-brown, densely pale yellow hairy) and abaxial leaf surface with domatia domed, with a central aperture (vs domatia absent or obscure), the secondary stem leaf bases acute (vs obtuse to truncate), the bracts forming a laciniate sheath on the distal peduncle (vs two opposite triangular bracts), the pyrene surface honeycombed with pits (vs entire).

## DISTRIBUTION & HABITAT

Lowland evergreen forest, rarely submontane. It has also been recorded once in secondary forest, and once in secondary thicket. Alt. range 15 – 960 m.

## CONSERVATION STATUS

*Keetia magassoubiana* has a wide geographic range but within this range it is infrequent and rarely collected, being recorded from a single specimen at most of the 14 recorded localities, and sites being widely separated. Its survival is threatened by clearance of its severely fragmented forest habitat. Over 90% of original forest was considered lost in Guinea by the end of the 20^th^ Century, similar to that of Sierra Leone (Sayer *et al*.1992). In Ivory Coast, 80 to 90% of forest surviving was lost in the period 1950 to 1980 (Riezebos *et al*. 1994). Historic sites for the species in Sierra Leone such as Kumrabai in Sierra Leone, where it was recorded in 1914 by N.W. Thomas (*N. W. Thomas* 7068, K!) are now entirely devoid of any of original forest habitat, which has been replaced by vast areas of oil palm cultivation (observed on Google Earth Pro, 15 June 2024).

At the eastern end of the range, the Adiopodoumé Forest in Ivory Coast, from which this species was recorded in the 1950 to 1970s, it is also likely extinct. Here its forest habitat has been lost due to anthropogenic disturbances such as the felling of trees for commercial purposes, land clearing for agriculture and uncontrolled urbanization which have led to the extinction of other species (Laurent 2020). The species is also extinct at its only known locality in the Sula Mts of Sierra Leone due to open-cast iron ore mining at the Tonkolili project managed since 2020 by the Kingho mining company (https://www.mining-technology.com/projects/tonkolili-iron-ore-mine/ accessed 15 June 2024). Recorded at Tonkolili in 2009 (*Robinson* 5) through an environmental impact assessment before mining began, the site of the specimen collection is now entirely denuded of vegetation (Google Earth Pro imagery observed June 2024).

However, the species is likely to survive at its site in the Tai Forest of Ivory Coast, which is considered well protected, and at Mt Bero in Guinea as a designated Important Plant Area (Couch *et al*. 2019,Darbyshire *et al*. 2017). In total it is thought to survive at five known sites, being lost from c. eight sites and with one site `sources du Niger’ not pinned down.

Here it is assessed as Endangered EN B1ab(iii) in view of the AOO of 20 km^2^, number of locations five and extreme human-induced habitat fragmentation and ongoing threats. One surviving site was recorded for an environmental impact assessment of a proposed mine, another two locations are entirely unprotected in a region where even in protected areas, virtually all natural habitat is at risk of clearance for development.

### COLLECTIONS STUDIED. GUINÉE (CONAKRY)

Gallery forest at the source of the Niger, (8°19’N 8°41’W), 330 m, fr. 1 Oct. 1944, *P. Jaeger* 140 (K! barcode K001900250); **Boké**, Boffa Préfecture, 10°25’23”N 14°19’07”W, 112 m, fr. 18 Oct. 2016, *E. Bidault* 2434 (K! barcode K000593387, MO); **Nzérékoré Préfecture**, between Lamineta village and Mont Kouoye, 8°12’50”N 8°37’53”W, 960 m, fl. 22 Nov. 2008, *F. Fofana* 170 (HNG, K! barcode K000615446, SERG, WAG); **SIERRA LEONE. Kukuna**, (9°8’28.4”N 12°39’52.1”W, 56m), fl. 17 Jan. 1891, *Scott Elliot* 4503 (K! barcode K000874710); **Kumrabai**, (8°41’23.9”N 13°7’54.3”W, 15 m), 31 Dec. 1914, *N. W. Thomas* 7068 (K! barcode K000874711); (8°41’23.9”N 13°07’54.3”W), 1 Dec. 1914, *N. W. Thomas* 6944 (K! barcode K000874712);**Tonkolili**, Northern region, 8°58’11.8”N 11°41’30.8”W, 770 m, fl. 6 Dec. 2009, *E. Robinson* 5 (FBC iso., K holo! barcode K000678056, SL iso.); Northern region, (8°58’51.6”N 11°47’52.8”W), 128 m, fl. 1 Dec. 1950, *E. L. King*, 199 (K! barcode K000593371); **LIBERIA. Nimba**, Grassfield, 7°29’N 8°34’W, 560 m, fr. 16 May 1970, *J. G. Adam* 25533 (K! barcode K000874719, P! barcode P03938759); Mt Nimba, Forest edge area, 7°32’N 8°33’W, 500 m, fl. 15 Dec. 1966, *J. J. Bos* 2436 (WAG! barcode WAG1312264, K! barcode K000874722); **Lofa**, Gbarnga-Zorzor Road, West bank of St. Paul, 7°19’N 9°30’W, 285 m, 20 Dec. 1966, *J. J. Bos* 2489 (WAG! Barcode WAG1312262, K! barcode K000874720); **IVORYCOAST. Bouna**, National Park North side, 9°36’N 3°40’W, 637 m, fr. 24 Aug. 1963, *W. J. J. O. de Wilde* 762 (P! barcode P03983357, WAG! barcode WAG1312258, BR! barcode BR0000020283841, K!); **Oroumbo-Bocca**, near Assakra, 10°36’N 14°20’W, 400 m, 7 Nov. 1961, *J. J. F. E. de Wilde* 3225 (WAG, K!); **Adiopodoumé**, Adiopodoumé Forest, (5°21’N 4°08’W), 60 m, 11 Feb. 1958, *Aké Assi* 5321 (K! barcode K000874726); Adiopodoumé Forest, (5°21’N 4°08’W), 60 m, 12 Dec. 1959, *Aké Assi* 5454 (K! barcode K000874727);Adiopodoumé Forest, (5°19’53”N 4°08’02”W), 40 m, 17 May 1964, *Leeuwenberg* 4191 (WAG, K! barcode K000874729); **Soubré**, Taï National Park, (5°46’N 6°54’W, 195 m), 18 Dec. 1975, *Aké Assi* 13154 (K! barcode K000874728).

### ETYMOLOGY

Named for Dr Sekou Magassouba, Director-General of the National Herbarium of Guinea (HNG) in the University of Gamal Abdel Nasser – Conakry, Republic of Guinea. Under his careful, tireless and diligent administration, HNG has increased greatly in its capacity to devise and manage projects, attract grants, to train students, including now at doctorate level for the first time, and to develop publications to research and publicise the conservation of the threatened plant species and habitats of his country.

### PHENOLOGY

Flowering late Nov. and Dec. Fruiting Feb.-May.

### VERNACULAR NAMES AND USES

None are known.

**Key to Species of *Keetia* in Upper Guinea (Africa West of the Dahomey gap)**

1. Stems with long (> 1.5 mm), erect, red or black hairs on the internode below the apex; leaf bases cordate on principal axis ................................. **14**

1.Stems glabrous or, if hairs present, adpressed and/or < 0.5 mm long, usually white; leaf bases of principal axis rounded to cuneate ........................... **2**

2.Tertiary nerves strongly scalariform, conspicuous, closely spread ............. **3**

2.Tertiary nerves inconspicuous or, if conspicuous not scalariform but reticulate ....... **4**

3.Leaves small, < 3 cm wide; stems densely red hairy .............. *K. bridsoniae*

3.Leaves large, > 3 cm wide; stems glabrescent ................... *K. venosa*

4.Leaf-blades obovate, apex truncate ......................... *K. obovata*

4.Leaf-blades ovate-elliptic or elliptic oblong, apex rounded, acute, obtuse or acuminate .. **5**

5.Stems glabrous (first internode below apex) .......................... **9**

5.Stems with hairs (first internode below apex), < 0.5 mm long, either adpressed and/or erect and (and then usually dense) ................................... **6**

6.Stems with hairs sparse (sometimes absent) red, thick, > 1.5 mm long, adpressed, (first internode below apex); tertiary nerves inconspicuous. Domatia glabrous, dome-like .................................. *K. magassoubiana sp. nov*.

6.Stems with hairs yellow, red or white, erect, < 0.5 mm long, usually dense. Tertiary nerves conspicuous (except *K. tenuiflora*). Domatia hairy, flat and open ................ **7**

7.Leaves drying black; tertiary nerves not conspicuous; stem hairs dense, yellow …..........................................*K. tenuiflora*

7.Leaves drying grey-green or dull red, tertiary nerves conspicuous; stem hairs white or red . ..................................................**8**

8.Stem hairs white; quaternary nerves inconspicuous, reticulum absent; leaf-blade < 6 cm long, usually densely hairy on lower surface ........................ *K. cornelia*

8.Stem hairs red; quaternary nerves forming conspicuous reticulum with tertiary nerves; leaf-blade 6 – 12 cm long, glabrous or sparingly hairy on lower surface ....... *K. venosissima*

9.Stems and leaves drying red-brown; tertiary and quaternary nerves conspicuous; stipules ovate-elliptic c. 11 x 8 mm .............................. *K. rubens*

9.Stems and leaves drying black or green; quaternary nerves inconspicuous; stipules with a long and slender apex, more than twice as long as broad .................... **10**

10.Fruit (fully-formed) ripening yellow-orange, 9 – 11 mm wide, disc not accrescent (< 2mm diameter), domatia with erect tufts of hairs or pit-like, glabrous ................ **11**

10.Fruit (fully-formed) ripening green or black, 14 – 25 mm wide, disc massively accrescent (> 4 mm diameter), domatia broad, flat, with horizontal red hairs or domed with apical minute aperture and glabrous .....................................**12**

11.Domatia with erect tufts of hairs, flat; stipule apex keel-like (laterally flattened); tertiary nerves often inconspicuous; midrib orange ...................... *K. mannii*

11.Domatia pit-like, glabrous; stipule apex rounded in cross-section; tertiary nerves always conspicuous; midrib concolorous, green ...................... *K. multiflora*

12.Fruit ripening black, 14 – 20 mm wide, disc massively accrescent (> 4 mm diameter), domatia broad, flat, with horizontal red hairs .......................... **13**

12.Fruit ripening green, 20 – 25 mm wide, disc massively accrescent (> 4 mm diameter), domatia domed with apical minute aperture and glabrous ..... *K. magassoubiana sp. nov*.

13.Leaves drying black; pedicels glabrous in young fruits, spur branches (brachyblasts) erect; fruit base rounded .............................. *K. abouabou*

13.Leaves drying green; pedicels hairy in young fruits; spur branches (brachyblasts) reflexed; fruit base cordate

................................. *K. susu*

14.Principal axis inflated at intervals, hollow, inhabited by ants; stem and leaves drying black ........................................... *K. hispida*

14.Principal axis lacking swellings, solid in section, not inhabited by ants; stems and leaves drying grey-green, or red ....................................**15**

15.Leaves and stems drying red-brown; leaves < 3 cm wide ............ *K. leucantha*

15.Leaves and stems drying grey-green, not red; leaves > 3.5 cm wide ............**16**

16.Stems densely hairy; hypanthium with dense ruff of white hairs; stipule arista longer than wider basal part .................................... *rufivillosa*

16. Stems very sparsely red hairy; hypanthium glabrous; stipule ovate-oblong, arista absent .. …..............................................*K. futa*

### NOTES

*Keetia magassoubiana* is recognised by its relatively small, seemingly glabrous narrowly elliptic-oblong leaves which dry black, as do the stems. Under the microscope the petioles and lower surface of the leaf midribs can be seen to have sparse, thick, adpressed, reddish orange hairs c. 0.5 mm long. In some specimens these hairs extend to the first internode of the stem. The domatia, sporadic in the axils of the midrib and secondary nerves and bifurcations, are unusual being domed, glabrous, with a minute apical aperture. The secondary nerves proceed to the margin in a straight line until they abruptly fork, or angle upward to the nerve above, from occasional domatia.

The fully formed fruits are massive, c. 2.2 – 2.5 cm broad, glossy and deeply bilobed as are those of *K. abouabou* Cheek (a local coast endemic of Abidjan, possibly extinct) and *K. susu* Cheek (endemic to Littoral, Guinea). However, the fruits of *K. magassoubiana* differ from these in that the two carpels are more deeply separated (didymous), so that the fruit is notched at both base and apex, and the accrescent disc is V-shaped, since from the central style base, the two flanks spread up towards the apex of the two lobes. In *K. abouabou* and *K. susu*, the two carpels are not separated from each other at the base, and the accrescent disc is not V-shaped, but spreads in a single plane.

*K. magassoubiana* can be separated from *K. multiflora* with which it has been confused in FWTA (Hepper 1963) and from *K. tenuiflora* to which it appears most closely similar (Bridson 1986) using the characters in Table 1.

Bridson annotated many of the specimens at K that are cited here as “sp. nov. aff. tenuiflora”.

About 75% of new plant species to science published today are already threatened (Brown *et al*. 2023). Usually this is because they have small ranges making them at risk of extinction from habitat clearance (Cheek *et al*. 2020b). *Keetia magassoubiana* is unusual in having a wide range yet is still massively threatened due to the extent of habitat clearance. Conservation actions are needed if it is not to become globally extinct as have so many other plant species (Humphreys *et al*. 2019;Cheek *et al*. 2018b)

## Supporting information

Supplemental file Holotype image

## Declarations

The authors declare that they have no conflicts of interest.

## Acknowledgements

Shigeo Yasuda, Charles Gore, and Amy Guest, assisted with specimen measurements and extracting pyrenes from fruits. Emily Robinson (now Harris) of SRK collected the type specimen, Iain Darbyshire made the photos of the new species in flower at Tonkolili and gave permission to use them.

Julie Edwards is thanked for support of the Sierra Leone TIPAs project and we thank Franklinia Foundation and JRS Biodiversity for supporting our field and plant species conservation work in Guinea.

