## Supplemental file Holotype image for "*Keetia magassoubiana sp*. *nov*. (Rubiaceae - Vanguerieae) a threatened evergreen forest climber of West Africa"

HERB. HORT. KEW.

ROYAL BOTANIC GARDENS KEW
K000678056

~~Keetia darwiniana ined. in Cheek et al 2017/2018~~

DET. M. Cheek 2 viii 2017

Holotype of Robinson 5

Keetia magassoubiana Cheek ined.

DET. CHEEK 20 June 2024

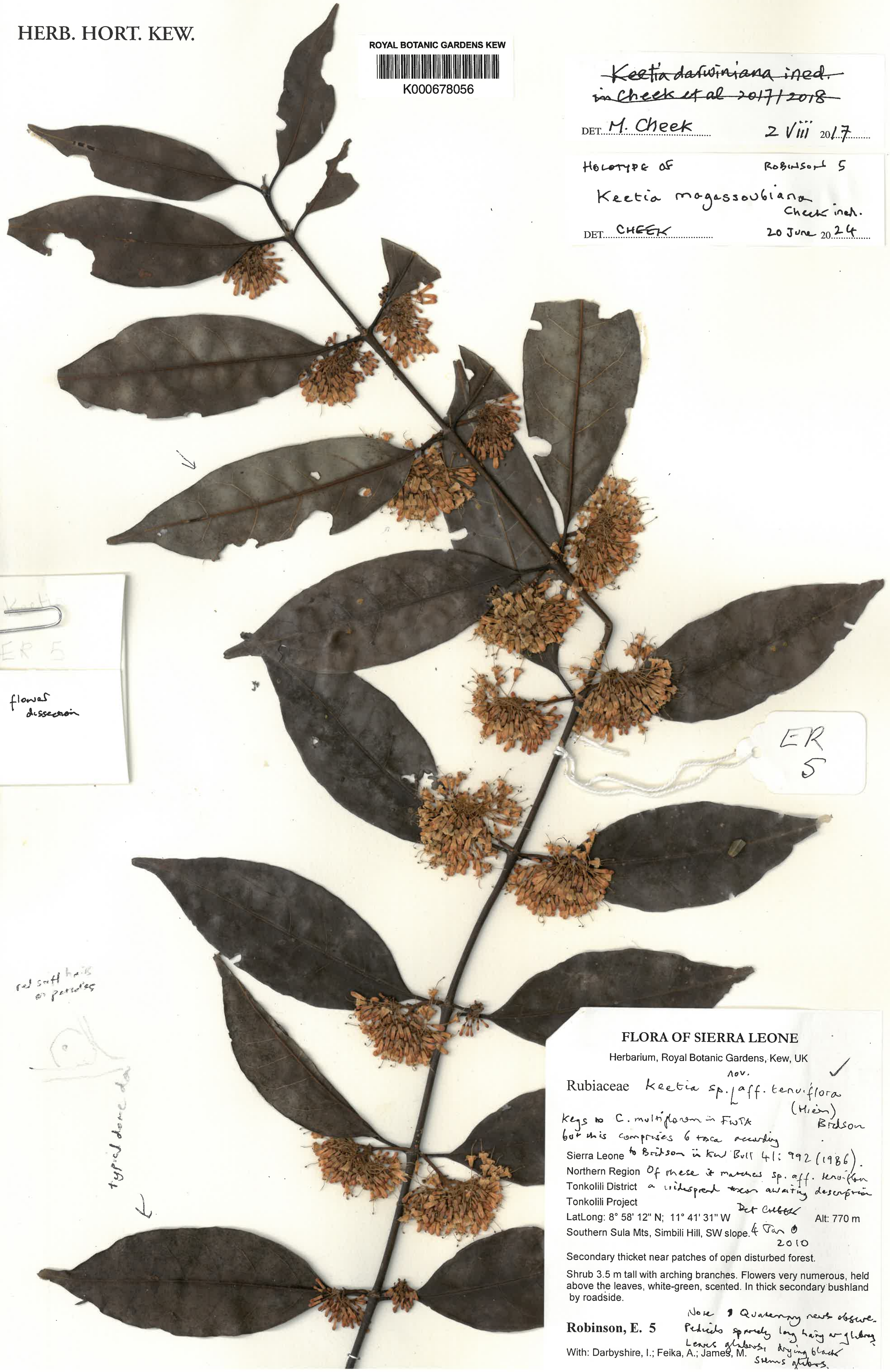

ER 5

flower dissection

ER 5

red soft hairs on petioles

typical dome da

### FLORA OF SIERRA LEONE

Herbarium, Royal Botanic Gardens, Kew, UK

Rubiaceae Keetia sp. nov. [aff. tenuiflora (Hiern) Bridson

Keys to C. multiflorum in FWTA but this comprises 6 taxa according to Bridson in Kew Bull 41: 992 (1986). Of these it matches sp. aff. tenuiflora a widespread taxon awaiting description Det Cheek 4 Jan 2010

Sierra Leone
Northern Region
Tonkolili District
Tonkolili Project
LatLong: 8° 58' 12" N; 11° 41' 31" W Alt: 770 m
Southern Sula Mts, Simbili Hill, SW slope.

Secondary thicket near patches of open disturbed forest.

Shrub 3.5 m tall with arching branches. Flowers very numerous, held above the leaves, white-green, scented. In thick secondary bushland by roadside.

Note: Quaternary nerves obscure. Pedicels sparsely long hairy or glabrous. Leaves glabrous, drying black. Stems glabrous.

**Robinson, E. 5**

With: Darbyshire, I.; Feika, A.; James, M.
